# Newly developed structure-based methods do not outperform standard sequence-based methods for large-scale phylogenomics

**DOI:** 10.1101/2024.08.02.606352

**Authors:** Giacomo Mutti, Eduard Ocaña-Pallarès, Toni Gabaldón

## Abstract

Recent developments in protein structure prediction have allowed the use of this previously limited source of information at genome-wide scales. It has been proposed that the use of structural information may offer advantages over sequences in phylogenetic reconstruction, due to their slower rate of evolution and direct correlation to function. Here, we examined how recently developed methods for structure-based homology search and tree reconstruction compare to current state-of-the-art sequence-based methods in reconstructing genome-wide collections of gene phylogenies (i.e. phylomes). While structure-based methods can be useful in specific scenarios, we found that their current performance does not justify using the newly developed structured-based methods as a default choice in large-scale phylogenetic studies. On the one hand, the best performing sequence-based tree reconstruction methods still outperform structure-based methods for this task. On the other hand, structure-based homology detection methods provide larger lists of candidate homologs, as previously reported. However, this comes at the expense of missing hits identified by sequence-based methods, as well as providing homolog candidate sets with higher fractions of false positives. These insights help guide the use of structural data in comparative genomics and highlight the need to continue improving structure-based approaches. Our pipeline is fully reproducible and has been implemented in a snakemake workflow. This will facilitate a continuous assessment of future improvements of structure-based tools in the Alphafold era.

## Introduction

As the number of available genomes increases, so does our ability to investigate deeper and unresolved evolutionary questions. However, traditional phylogenetic methods based on sequences have limitations, such as the saturation of the phylogenetic signal when dealing with highly divergent sequences and/or distantly related species. When seeking for new sources of phylogenetic information, protein structures have been considered based on the idea that they are expected to diverge at a slower pace than sequences (Illergård et al. 2009). However, the scarcity of structural information has limited their use in large-scale phylogenetic studies.

The development of AlphaFold2 has recently revolutionised the field of structural biology, unlocking the analyses of protein structures at scales previously considered unachievable (Jumper et al. 2021). As a consequence, new methods have been developed to handle such scales. One example is Foldseek: a software that allows fast alignment of protein structures based on a recoding of the structural information into a 20-states alphabet (Van Kempen et al. 2024). Recently, Moi et al. 2023 developed Foldtree: a pipeline to quickly compute distance-based phylogenies from protein structures which the authors argued can outperform traditional methods also for closely related proteins.

Motivated by the potential shown by Foldtree, we analysed whether a phylogenomic pipeline could be adapted to use protein structures for genome-wide phylogenetic analysis, such as the reconstruction of all evolutionary histories of human genes -i.e-the human phylome (Huerta-Cepas et al. 2007).

## A pipeline to benchmark structure versus sequence phylomes

A set of 18 eukaryotic species (Tab. S1) was selected to reconstruct the human phylome based on available proteomes from UniProt (The UniProt Consortium et al. 2023). By inferring the human phylome, we refer to reconstructing the phylogenetic relationships between all *H. sapiens*’ proteins (seed/query proteins) and their homologs in the species set (target proteins).

Protein structures were downloaded from the AlphaFold database (Varadi et al. 2024). Proteins with poorly predicted structures (average predicted Local Difference Distance Test (pLDDT) < 40) were excluded. Protein sequences were extracted from the structure files to ensure full congruence between sequences and structures (Fig. 1A).

**Figure 1.**
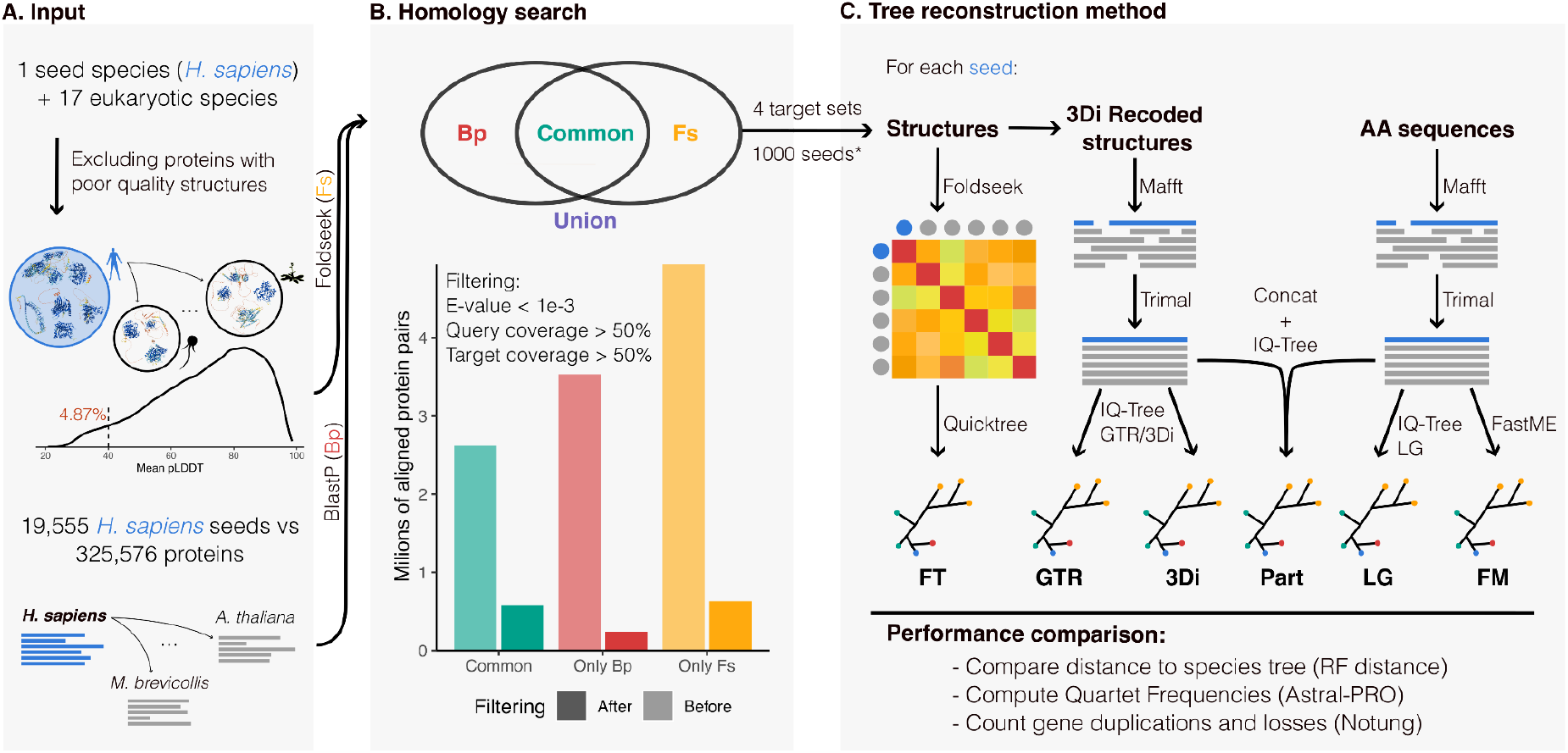
Schematic representation of the pipeline. A) Primary amino acid sequences and 3Di-recoded structures from *Homo sapiens*’ proteins (seeds) are aligned against a dataset of 18 eukaryotic species with BlastP (*Bp*) and Foldseek (*Fs*), respectively. B) Before entering into the phylogenetic pipeline, *Bp* and *Fs* results are divided into four target sets per seed as shown in the Venn diagram. The number of query-targets pairs in each target set before and after filtering is shown below. C) The four target sets of each seed are submitted to six tree reconstruction methods. (For computational reasons, we restricted step C to 1000 randomly selected seeds. *Among the randomly selected seeds, only those with at least four common hits entered into step C.

The sequence and the structure of each *H. sapiens* seed protein was aligned with BlastP and Foldseek, respectively, against the 18 species’ proteomes dataset (Altschul et al. 1990; Van Kempen et al. 2024). Both results were filtered based on E-value < 1e-3, query coverage > 50%, and target coverage > 50%. Four target sets were retrieved for each seed: top 150 BlastP hits (*Bp*), top 150 foldseek hits (*Fs*), *Union*, and intersection (*Common*) between *Bp* and *Fs* sets (Fig. 1B).

For each target set, six tree reconstruction methods were applied (Fig. 1C and supplementary methods for details):

1. *FT*: structure-based distance method. The same procedure as the FoldTree snakemake implementation (Moi et al. 2023).
2. *3Di*: structure-based Maximum Likelihood (ML) method. Structures are recoded with the 3Di alphabet computed by Foldseek. After masking sites with pLDDT < 50, 3Di sequences were aligned with FoldMason (Gilchrist et al. 2024). The alignments were then trimmed with TrimAl -gappyout (Capella-Gutiérrez et al. 2009). Trees were reconstructed with IQ-Tree2 (Minh et al. 2020) specifying as substitution matrix the 3Di matrix computed in Puente-Lelievre et al. 2024.
3. *GTR*: structure-based ML method. The same alignment as *3Di* was used in IQ-Tree2 using the GTR substitution matrix (GTR20).
4. *LG*: sequence-based ML method. Protein sequences were aligned with mafft --auto (Katoh and Standley 2013) and trimmed as in *3Di* and *GTR*. Trees were reconstructed with IQ-Tree2 specifying the LG substitution matrix (Le and Gascuel 2008).
5. *FM*: sequence-based distance method. The same alignment for *LG* was used as input to FastME (Lefort et al. 2015).
6. *Part*: the trimmed alignments from *LG* and 3Di were concatenated and used as input in IQ-Tree using a partition scheme.

For computational reasons we subsampled 1000 random seeds, filtered those with less than 4 *common* hits, and proceeded with inferring 24 phylogenies per seed (four target sets per six tree reconstruction methods). We compared the accuracy of each gene tree reconstruction method according to three different metrics (see Supplementary methods for details).

All the scripts used in this project were developed as a reproducible and customisable pipeline in Snakemake (Mölder et al. 2021) available at https://github.com/Gabaldonlab/structural_phylome. These analyses can be reproduced with any taxon set present in UniProt. Check the Supplementary Methods for additional information.

## Divergences between BlastP and Foldseek

Among the query-target pairs identified by BlastP and Foldseek, only 2.62*10^6^ pairs (23.6%) were detected by both tools (*common* targets). After filtering, this proportion increases to 39.8% (0.58*10^6^ pairs), highlighting a significant disparity between the two approaches (Fig. 1B). The two tools respectively identified 3.53*10^6^ (31.8%) and 4.94*10^6^ (44.5%) unique hits (referred to as *singletons* hereafter). These proportions expectedly decrease to 16.6% and 43.6% after filtering.

To understand these striking differences, we investigated three alignment metrics in the unfiltered results: percentage of identity, E-value and query coverage. For each metric we explored the correlation among BlastP and Foldseek in *common* hits (see binned scatterplots in Fig. 2A-C), and the distribution of values in *singletons* and *common* hits for these metrics (see marginal distributions in Fig. 2A-C). We observed that the percentage of identity of *common* hits is highly correlated between BlastP and Foldseek (r=0.85, Fig. 2A). Interestingly, only Foldseek *singletons* have lower identity values (mode < 25%) when compared to *common* hits (Fig. 2A). This indicates that Foldseek, as proposed, may be useful to detect highly-divergent homologs in the so-called sequence identity “twilight zone” (Sander and Schneider 1991; Rost 1999; Puente-Lelievre et al. 2024). However, in an assessment based on the EggNOG orthologous groups (Huerta-Cepas et al. 2019), we found that the potential of Foldseek in identifying remote homologs comes at the expense of missing a higher fraction of established co-orthologous relationships when compared to BlastP (Fig. S1A) The E-values of both tools show moderate correlation (r=0.43) but Foldseek E-values tend to be lower (Fig. 2B). BlastP and Foldseek *singletons* have similar E-value distributions when compared to *common* hits. Regarding query coverage, BlastP singletons usually have lower values for this metric (see red marginal distribution in Fig. 2C). Consistent with this, the hits identified by BlastP but not by Foldseek in the EggNOG orthologous groups analysis usually show low query coverage values (Sup Fig. S1B). This suggests Foldseek may be missing hits when homology conservation is restricted to a smaller fraction of the protein space.

**Figure 2.**
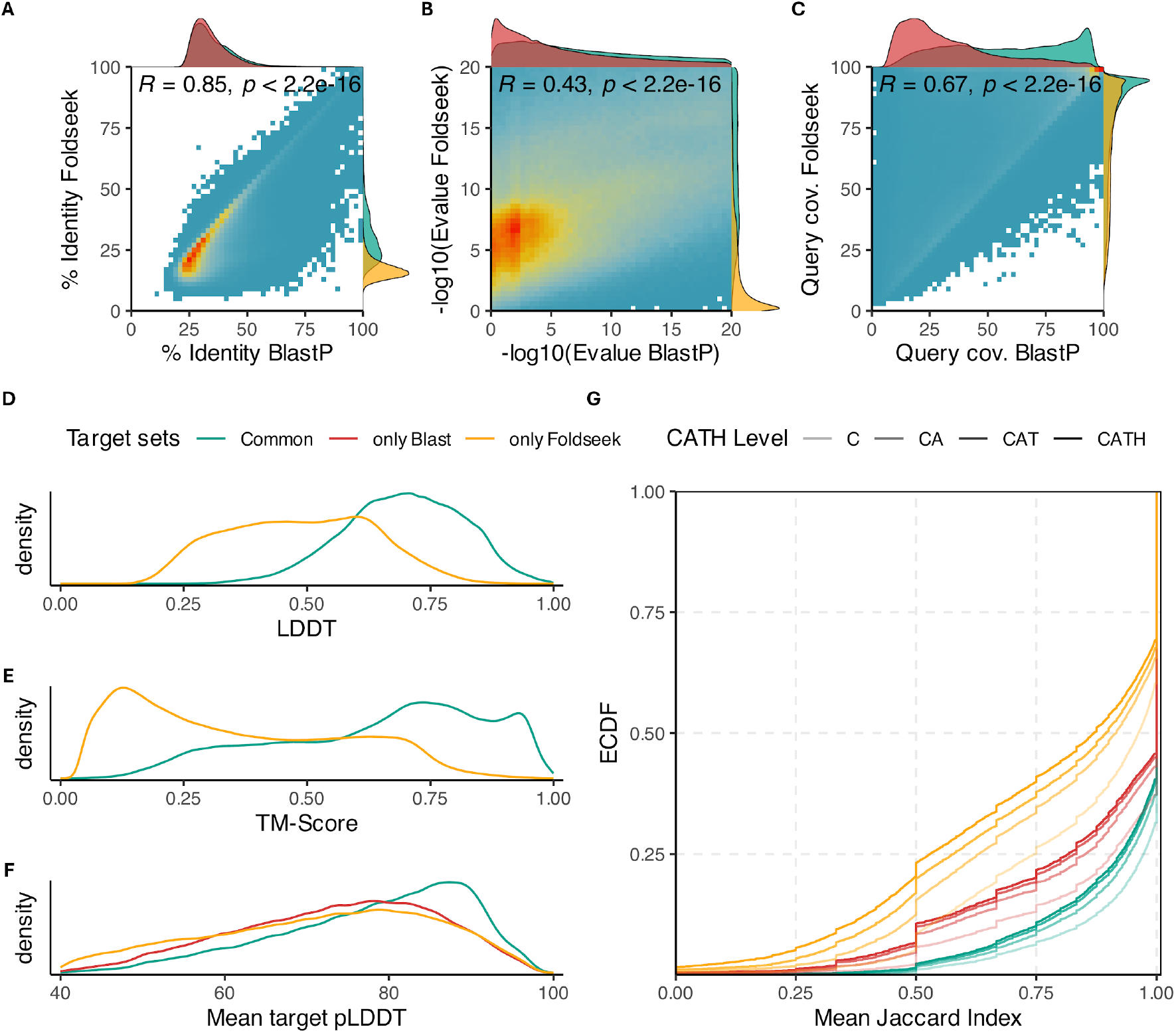
Distributions and correlation of A) percentage identity, B) -Log10(E-value) and C) query coverage for unfiltered BlastP and Foldseek results. The binned scatter plot colour scale goes from light blue (low density) to red (high density). The marginal distributions are colour-coded according to the target set. Distribution of D) Local Difference Distance Test (LDDT), E) Template-Modelling score (TM-score) and F) mean target predicted LDDT between different target sets. G) Cumulative distribution of mean Jaccard index per query for all levels of CATH annotation, including protein class (C), architecture (A), topology (T) and homologous superfamily (H) (coded with different transparency levels, see legend). See Supplementary Methods for details on this analysis.

For each alignment, Foldseek computes both the average Local Difference Distance Test (LDDT) and the Template Modelling Score (TM). These two metrics quantify the overlap between pairs of aligned structures and both are clearly lower in Foldseek *singletons* indicating highly divergent homologs or possible spurious homologs (Fig. 2D-E). Regarding the quality of protein structure predictions, *common* hits have higher average pLDDT (as computed by AlphaFold) compared to both Foldseek and BlastP *singletons* (Fig. 2F). This indicates that poorly predicted structures may be responsible for both false positives (Foldseek *singletons*) and false negatives (BlastP *singletons*) in Foldseek searches.

As an orthogonal benchmark of the two similarity search methods we explored the homogeneity of CATH annotations in filtered BlastP and Foldseek results. Our rationale was that correct homologous identifications should result in target sets with higher fractions of shared C, CA, CAT and CATH domain annotations with respect to the query. *Common* hits have more homogenous annotation, followed by BlastP *singletons* (Fig. 2G). The lower levels of concordance of Foldseek *singletons* suggests that this method recovers a higher fraction of non-homologous targets than BlastP.

## Underperformance of structure-based methods for tree reconstruction

We compared the performance of sequence-based vs structure-based tree reconstruction methods based on the following metrics (Fig. 3): (1) Robinson-Foulds (Robinson and Foulds 1981) distance to the species tree of single copy orthologs subtrees obtained from splitting multicopy gene families using DISCO; (2) fraction of gene tree quartets agreeing with the species tree topology (quartet support), and (3) number of duplication and losses needed to reconcile the gene trees with the species tree. More accurate gene trees are expected to have lower RF distances to the species tree, show higher quartet support values, and require less events to be reconciled.

**Figure 3.**
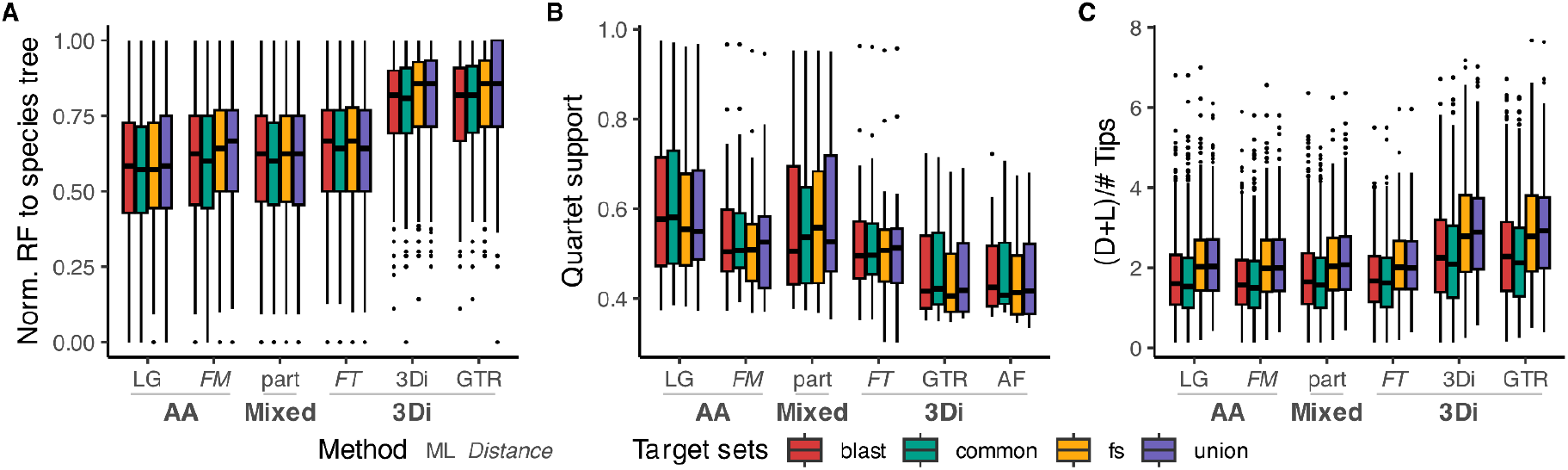
Boxplot distribution grouped by target set and tree reconstruction method of A) normalised Robinson-Foulds (RF) distance of decomposed single copy gene trees to the species tree, B) First Quartet Frequency support values (this measure indicates how many times the nodes in the species trees are observed in the gene trees), and C) number of gene duplications and losses inferred by gene tree-species tree reconciliation normalised by number of tips.

Structure-based methods (*3DI, GTR, FT*) did not outperform sequence methods (*LG, QT*) in any of these metrics (Fig. 3). All the ML structure-based methods (*3DI, GTR*) clearly underperformed the ML sequence-based method (*LG*). Bootstrap support values were higher in *LG*, perhaps due to the worse performance of ML structure-based methods (Fig S2A). Both distance methods, sequence-based (*FM*) and structure-based (*FT*), performed similarly (Fig. 3). Further, they also produced very similar trees to each other (Fig. S2B). This suggests that structure and sequence may provide similar phylogenetic information but that structural ML methods still have room for improvement.

*LG* outperformed structure-based and sequence-based distance methods, most clearly in metrics (1) and (2) (Fig. 3A-B), inferring less discordant trees with the species tree for all target sets. This implies that *LG* performed better also for the target sets which include Foldseek *singletons* (i.e. candidate homologs only identified based on structural information). The better performance of ML sequence-based was confirmed in an independent benchmarking done with one of the datasets that were used to benchmark FoldTree (Moi et al. 2023) (Fig. S3, see Supplementary Methods). Finally, we also explored a partition scheme combining sequence and 3Di information (*part*), as done in a recent study (Puente-Lelievre et al. 2024). Interestingly, this condition outperformed ML structure-based methods, but does not seem to improve the performance of the ML sequence-only condition (*LG*) despite being more computationally intensive and time consuming.

To complement the previous analyses done on the distinct target sets, we explored the topological information of the gene trees. Foldseek *singletons* show a greater normalised internode distance to the seed compared to BlastP *singletons*, median distance of 0.625 and 0.547, respectively, based on *LG*, the best performing tree reconstruction method (this result is consistent in all tested conditions, Fig. S2C). *Common* hits were the closest to the seed (median=0.455), showing a more similar distribution to BlastP *singletons* than to Foldseek *singletons* (Fig. S2C). Overall, these results are concordant with Foldseek being able to detect homologs in the “twilight zone”. However, this comes at the expense of identifying a higher fraction of targets with non-homologous structures than BlastP (Fig. 2G), and also of missing BlastP *singletons*, which tend to be phylogenetically closer to the seed than Foldseek *singletons* (Fig. S2C).

As done in Moi et al. 2023, we assessed ultrametricity (measured as the variance of root-to-tip distances) as a possible index of tree reconstruction quality. We noticed that *FT* was considerably more ultrametric than any other method, as reported in Moi et al. 2023 (Fig. S2D). The two ML structure-based methods (*3Di* and *GTR*) were more ultrametric than *LG*, possibly indicating that structures may in fact present a more “molecular clock-like” behaviour than sequences. Notwithstanding this, since *LG* outperformed all structure-only based methods (Fig. 3), this indicates that ultrametricity is not necessarily an indicator of accurate phylogenetic reconstruction.

## Conclusions

Phylogenomic analyses based on structural information are now possible thanks to tools such as Foldseek and Foldtree. Despite this being a very important achievement, our results altogether point that structure-based methods still do not outperform the standard sequence-based tools for large-scale phylogenetics, particularly in the phylogenetic reconstruction step. The potential of the structure-based tool Foldseek to detect highly-divergent homologs in the “twilight zone” (Fig. 2A) could be of particular interest when protein sequences provide limited resolution for homology search (Himmel et al. 2023; Köstlbacher et al. 2024). However, its usage comes with the risk of missing sequences detected as homologs by Blast (BlastP *singletons*) which are more phylogenetically proximal to the seed sequence than Foldseek *singletons* (Figs. 2G and S2C). Regarding gene tree reconstruction, the ML sequence-based method performed better than *FT*, even when run on the set of structural homologs identified by Foldseek (Fig. 3). While ML sequence-based performed much better than ML structure-based for tree reconstruction, little differences in performance were observed between sequence and structure-based tree distance methods (Fig. 3). This suggests that future improvements either in the recodification of structural-information (Edgar 2024) and/or in ML implementations of structure-based tree reconstruction methods (Puente-Lelievre et al. 2024) could eventually lead structure-based methods to surpass sequence-based methods in large-scale phylogenomics.

## Data and Code Availability

All the data is already publicly available. The code used in this project, including how to recreate the same dataset and figures, is available at https://github.com/Gabaldonlab/structural_phylome.

## Supplementary Figures

**Figure S1.**
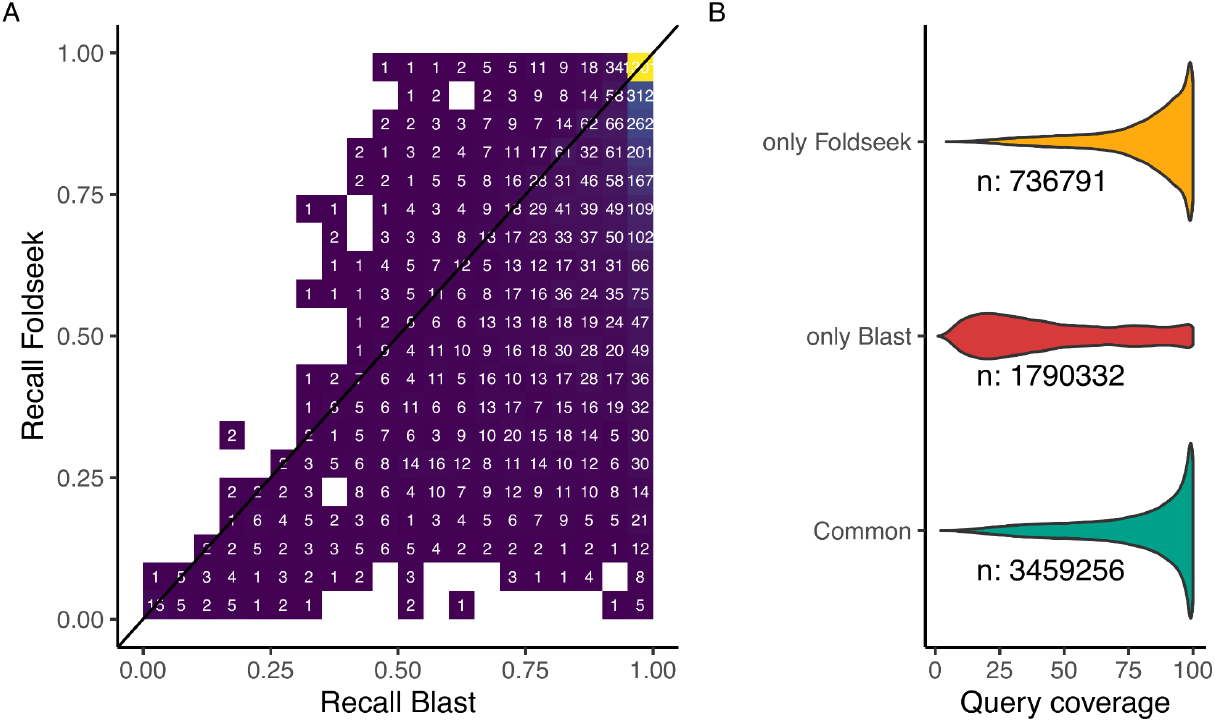
A) Heatmap comparing the fraction of pairs of proteins belonging to the same EggNOG group detected by Foldseek and by Blast. Numbers and color code indicate the density of data in each area of the plot. B) Query coverage percentage distribution of alignments between EggNOG homologs found by both tools (‘Common’), ‘only Foldseek’ and ‘only Blast’.

**Figure S2.**
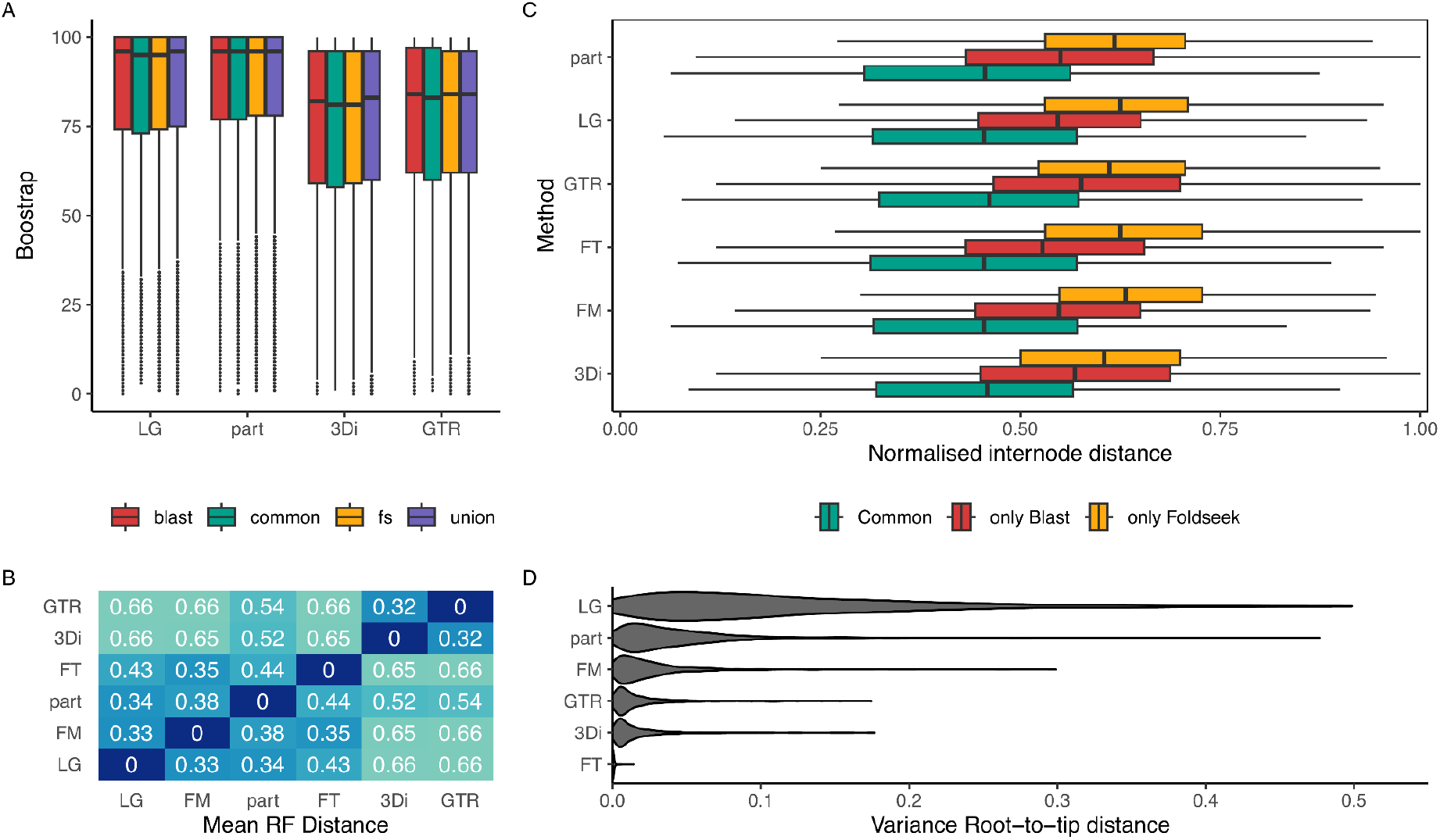
A) IQ-Tree ultrafast bootstrap values distribution binned by target set. B) Differences between tree reconstruction methods as measured by normalised Robinson-Foulds (RF) (median RF distance values are shown in the heatmap). C) Normalised internode distance to the seed (number of nodes from tip to seed divided by longest possible path to tip) distribution, coloured by target set and binned by tree reconstruction method in *union* trees. ‘Only Blast’ and ‘only Foldseek’ refer to BlastP *singletons* and to Foldseek *singletons*, respectively. D) Violin plot distribution of root-to-tip distances (in midpoint rooted trees) for common hits. Each violin is scaled in order to have the same maximum width.

**Figure S3.**
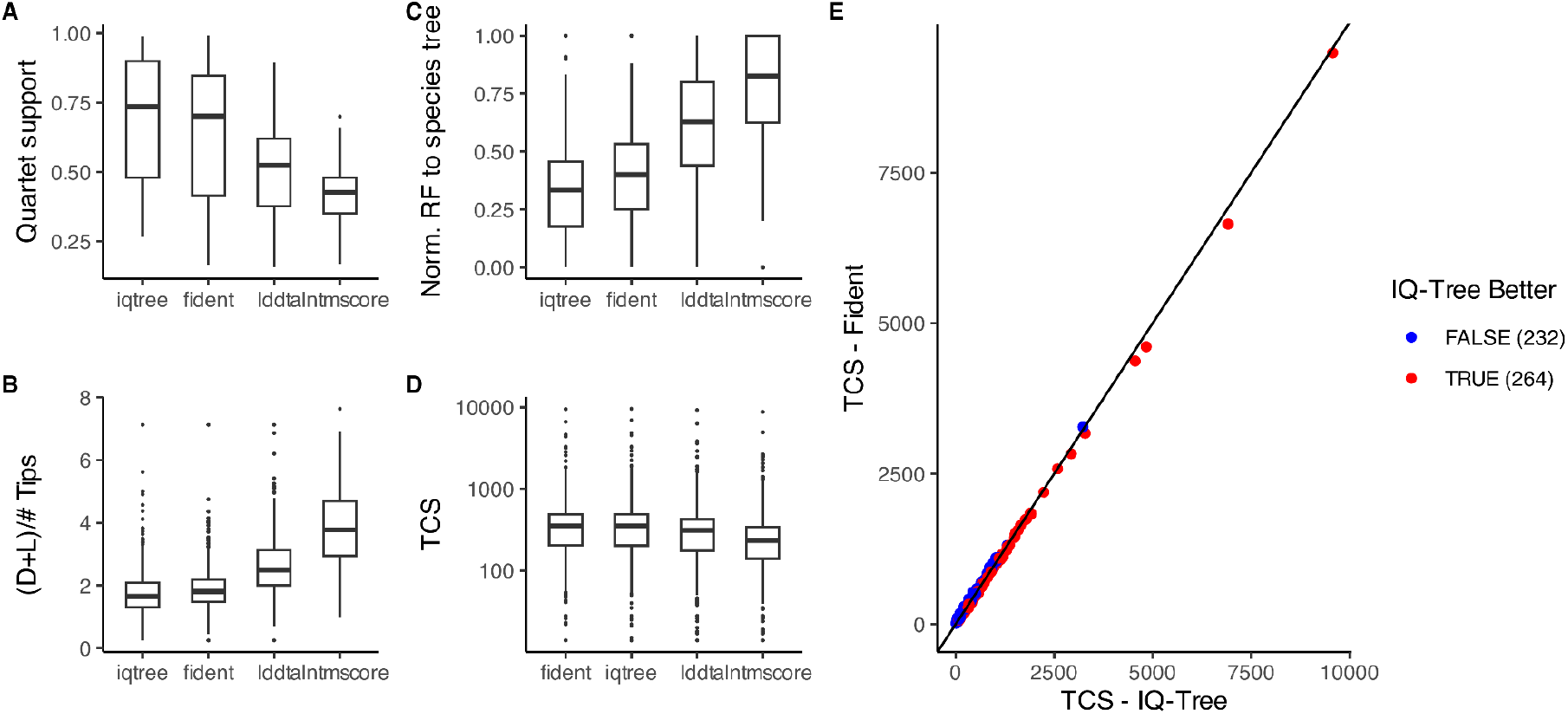
Boxplot distribution of A) First Quartet Frequency support values B) normalised Robinson-Foulds (RF) distance of decomposed single copy gene trees to the species tree, C) number of gene duplications and losses inferred by gene tree-species tree reconciliation normalised by number of tips, and D) Taxonomic Consensus score (TCS), divided by tree reconstruction method (‘iqtree’ represents a condition equivalent to LG; while ‘fident’ ‘lddt’ and ‘alntmscore’ are different metrics that were used by Moi et al. 2023 to run FoldTree). E) Scatterplot comparing the TCS values found by IQ-Tree and Foldtree (Fident), displayed in a similar fashion as done in Moi et al. 2023.

