## Supplementary methods for "Newly developed structure-based methods do not outperform standard sequence-based methods for large-scale phylogenomics"

### Material and methods

#### Data download

Structures are downloaded with gsutil v5.30 used to retrieve the cif.gz file for any given taxid present in UniProt. The files are then uncompressed and protein structures with mean pLDDT < 40 are excluded for downstream analyses. The protein sequences are directly extracted from the structures using the script cif2fasta.py from HH-suite v3.3.0 (Steinegger et al. 2019). This was done to have exactly the same set of proteins, which would not have necessarily been the case if we downloaded the proteomes directly.

#### Homology search

First of all we create a database for BlastP v2.16.0 (Altschul et al. 1990) and Foldseek v9.427df8a (van Kempen et al. 2024) with:

```
makeblastdb -in $seq -out $out_file -parse_seqids -taxid_map $taxidmap -dbtype prot
```

```
foldseek createdb $input $output
```

Then, each protein is searched with:

```
blastp -query $seed_proteome -max_hsps 1 -max_target_seqs 1000 -outfmt "6 std qcovs qcovhsp qlen slen staxids"
```

```
foldseek easy-search --max-seqs 1000 --format-output
query,target,fident,alnlen,mismatch,gapopen,qstart,qend,tstart,tend,
evalue,bits,lddt,alntmscore,rmsd,prob,qcov,tcov
```

Both Blast and Foldseek are filtered with E-value < 1e-3, query coverage and target coverage > 50%.

*Sequence retrieval:*
For 1000 non-orphan seed sequences we retrieve four different sets of homologs: top 150 foldseek hits, top 150 blast hits, common hits and union. If there are less than 4 common hits the seed is excluded from further analysis.

**EggNOG benchmark**
We obtained the eggNOG annotations for each protein in our dataset from UniProt. We retained the eggNOG groups containing more than five members and concatenated all the proteins associated with these selected orthologous groups. Using this concatenated file as both the query and target for BlastP and Foldseek using these commands:

```
makeblastdb -in {input.fa} -out {blast_db} -dbtype prot
```

```
blastp -query {input.fa} -db {blast_db} -out {blast} -max_hsps 1
-outfmt '6 std qcovs qcovhsp qlen slen' -max_target_seqs 500
```

```
foldseek createdb {input.tsv} {foldseek_db}
```

```
foldseek easy-search {input.db} {input.db} {foldseek}
$TMPDIR/eggnog_fs --max-seqs 500 --format-output
query,target,fident,alnlen,mismatch,gapopen,qstart,qend,tstart,tend,
evalue,bits,lddt,alntmscore,rmsd,prob,qcov,tcov
```

This approach allowed us to evaluate the fraction of members within the same eggNOG group aligning to each other according to each of the two tools. We took this metric as a measure of recall, since we would expect all members of the same eggNOG group to align to each other by the fact of being homologous (i.e., sharing similarity) to each other.

This benchmark can be reproduced with any phylogenomic database annotation available in UniProt using the `workflow/ortho_benchmark.smk` script available in the Github repository.

**Phylogenetic pipelines**
For each seed and target set six tree reconstruction methods are performed: (i) the recently proposed structure-based distance method Foldtree (*FT)* (Moi et al. 2023), two structure-based ML methods based on (ii) using a previously implemented 3Di-based substitution matrix (*3Di*) (Puente-Lelievre et al. 2024) or (iii) inferring a custom substitution matrix with GTR (*GTR*), (iv) a standard sequence-based Maximum Likelihood (ML) method using LG as substitution matrix (*LG*) (Le and Gascuel 2008), (v) a sequence-based distance method using FastME (Lefort et al. 2015) (*FM*) and finally a partitioned strategy mixing both aminoacid and recoded alignments (*part)*. Recoded refers to using the structures recoded with the 3Di alphabet. This alphabet has 20 letters and hence recoded structures can be processed downstream in the pipeline as if they were standard amino acid sequences. Therefore, the final trees will be 24 per seed sequence as the six pipelines will be applied to four different sequence sets. We also included the python implementation of Foldtree (*FT_PY*) that also includes core cutting and distance matrix correction. However, as the

current implementation is the Snakemake version and as the result did not significantly change we only kept *FT* in the manuscript (results not shown).

1. LG:

   Proteins are aligned with `mafft --auto` (v.v7.525) (Katoh and Standley 2013) and trimmed with `trimal -gappyout` (v1.5.0) (Capella-Gutiérrez et al. 2009). Then phylogenies are computed with IQ-Tree v2.3.6 with 1000 UltraFast bootstraps (Hoang et al. 2018; Minh et al. 2020):

   ```
   iqtree2 -s {input} --prefix $tree_prefix -B 1000 -T {threads}
   --boot-trees --quiet --mem 4G --cmin 4 --cmax 10 -mset LG
   ```

   This way heterogeneity rates and state frequencies are inferred with ModelFinder (Kalyaanamoorthy et al. 2017).

2. FT:

   In this case we followed the Snakemake implementation of Foldtree (https://github.com/DessimozLab/fold_tree) which includes first an all vs all foldseek search:

   ```
   foldseek easy-search $structdir $structdir {output} $TMPDIR
   --format-output 'query,target,fident,lddt,alntmscore'
   --exhaustive-search -e inf
   ```

   Then a pairwise identity matrix is obtained with `foldseekres2distmat_simple.py` which can then be used as input in quicktree as done in the snakemake implementation of Foldtree (Howe et al. 2002).

3. 3Di:

   The recoded structure sequences (in 3Di alphabet) are first masked (sites with pLDDT < 50 are annotated as “-”) . Then, `Foldmason` is used to align the recoded sequences with the 3Di Foldseek substitution matrix. The alignment is then trimmed with `trimal -gappyout`. The same IQ-Tree command as LG is applied but the model used comes from the 3Di rate matrix presented in Puente-Lelievre et al. 2024 and downloaded from https://github.com/nmatzke/3diphy.

4. GTR:

   The same pipeline as 3Di is applied but the GTR20 model is used in IQ-Tree (`--mset GTR20`).

5. FM:

   ```
   fastme -q -p -T {threads} -b {params} -i {input} -o {output} >
   {log}
   ```

   In this case FastME v2.1.6 is used (Lefort et al. 2015) after the fasta is converted to phylip with 100 bootstrap.

6. Part:

In this case the trimmed amino acid and 3Di alignments are concatenated. A partition file is created using as models the best LG and 3DI model (including rate heterogeneity and base frequencies parameters). IQ-tree is then with 1000 UltraFast bootstraps and the edge-proportional partition model:

```
iqtree2 -s {input.fa} -p {input.part} --prefix $tree_prefix
-mdef {params.threedi_submat} -B 1000 -T {threads}
```

**Performance assessment of tree reconstruction methods**

First we decompose multicopy gene trees in single copy trees with DISCO v1.4 (Willson et al. 2022). We then compute the Robinson-Foulds distance of these 1 to 1 ortholog gene trees with the species tree for those including at least more than 50% of our taxon sampling. We also compute the total number of duplications and losses as computed by Notung v.2.9.0, a reconciliation software based on parsimony (Stolzer et al. 2012). Finally, we run Astral-PRO v1.19.3.5 (Zhang et al. 2020), to quantify how many gene tree quartets support the species tree topology with:

```
astral-pro -c {input.sptree} -a {input.genemap} -u 2 -i {output.gt}
-o {output.st} -C --root {params}
```

**CATH benchmark**

CATH ('Homologous superfamily class', as annotated by Gene3D) domains are directly downloaded from UniProt thanks to the UniProt REST API (Lewis et al. 2018; The UniProt Consortium et al. 2023).

For each query-target pair, the Jaccard Index (JI) of the two sets of CATH IDs (the set from the query, and the set from the target) was computed for all levels of the database (C=class, A=architecture, T=topology/fold, H=homologous superfamily). We chose this metric to allow for the comparison of multidomain proteins.

For example, let's assume that for a given query-target comparison, query X is associated with the *1.10.8.270* domain, and target Y has two domain annotations: *1.10.8.270* and *1.10.10.750*. The JI for this query-target pair will be 0.5 at both *CATH* and *CAT* level. However, it will be 1 at *CA* and *C* level.

Only query-target pairs with CATH domain information were considered. After obtaining the JI for all pairs, we computed the mean value for each target class (*Common, only Blast, only Foldseek*) on a query basis.

**Benchmarking tree reconstruction methods with an alternative dataset**

We investigated how our results may differ to the ones reported in Moi et al. 2023. This was done by performing an independent analysis using their benchmarking pipeline (https://github.com/DessimozLab/fold_tree/blob/main/workflow/benchmarking) and our metrics (RF, quartet score and DL) in a set of 500 Eukaryotic OMA orthogroups (downloaded from https://doi.org/10.5281/zenodo.8346286) using the Quest for Orthologs eukaryotic species tree (available here: https://swisstree.vital-it.ch/ftp/speciestree.nhx and parsed in the get_sptrees.R script available in the GitHub repository). Summarising, for the *'iqtree'* mode we aligned sequences with `muscle` and then computed a tree with `iqtree -s $alignment -m LG+I+G -seed 42 -nt 1.` For *'fident' 'lddt'* and *'alntmscore'* we used Foldtree using the three distinct alignment properties to compute the distance matrix. We then computed RF distance, quartet support and DL events the same way as we did for the other trees. The Taxonomic Consensus Score (TCS) was computed using the script `calctreescores.py` included in the Foldtree repository. This benchmarking analysis is reproducible in the github repository and can be expanded to any orthogroups, following the instructions in the readme file.

**Reproducibility**

The pipeline can be reproduced with any taxon set available in UniProt just by installing Snakemake (developed in v8.11.6) (Mölder et al. 2021) and gsutil. All the rest of the software can be easily installed with Conda.

Each user may configure different filters including the minimum average pLDDT to exclude proteins (default 40), maximum number of targets in the tree (default 150) or Blast and Foldseek filters among others. The pipeline can be run in two modes: phylome and orthogroup. In the first case the user only needs to input a set of proteins present in the seed organism. Alternatively, the user can input a set of precomputed orthogroups.

The pipeline and more detailed documentation is available here: https://github.com/Gabaldonlab/structural_phylome.
